# Integrated clonal analysis reveals circulating tumor DNA in urine and plasma of glioma patients

**DOI:** 10.1101/758441

**Authors:** Florent Mouliere, Katrin Heider, Christopher G. Smith, Jing Su, Mareike Thompson, James Morris, Jonathan C.M. Wan, Dineika Chandrananda, James Hadfield, Marta Grezlak, Irena Hudecova, Wendy Cooper, Davina Gale, Matt Eldridge, Colin Watts, Kevin Brindle, Nitzan Rosenfeld, Richard Mair

## Abstract

Glioma-derived cell-free tumor DNA is challenging to detect using standard liquid biopsy techniques as its levels in body fluids are very low, similar to those in patients with early stage carcinomas. By sequencing cell-free DNA across thousands of clonal and private mutations identified individually in each patient’s tumor we detected tumor-derived DNA in plasma (10/12, 83%) and urine samples (8/11, 72%) from the majority (7/8, 87.5%) of glioma patients tested.

**One Sentence Summary:** Circulating tumor DNA can be detected in the majority of plasma and urine samples from primary brain tumor patients using sequencing guided by mutations detected in multi-region tumor biopsies.

## Introduction

Primary brain tumors, which are diagnosed in over 260,000 patients annually (1), have a poor prognosis and lack effective treatments. Better methods for early detection and identification of tumor recurrence may enable the development of novel treatment strategies including the use of immunotherapy (2, 3). The development of new treatments would also benefit from minimally invasive methods that characterise the glioma genome (4–6). DNA analysis in liquid biopsies has the potential to replace or supplement current ineffective imaging-based monitoring techniques and reduce the morbidity associated with repeated biopsy while providing the genomic information required by precision medicine (4, 7, 8). However, cell-free tumor DNA (ctDNA) is extremely challenging to detect in the plasma of patients with brain tumors as its fractional concentration (mutant allele fractions, MAF) are in the same range as those observed in plasma of patients with early stage carcinomas (9, 10). Reported detection rates for glioma are typically around 15%-30% (9), although higher rates of detection have been claimed (10). Tumor DNA has been detected in urine for some cancers, however this has been largely limited to urothelial cancers, or patients with advanced cancers having high tumor burden detected in plasma (11–13). Cerebrospinal fluid (CSF) has been proposed as an alternate medium for ctDNA analysis (14–18), however detection rates have been low as reported in a recent large study (42/85 patients detected, 49.4%) (19). In addition, CSF sampling via lumbar puncture is an invasive procedure, which limits its use for longitudinal sampling (20, 21).

Here we demonstrate that tumor DNA can be detected in the majority of plasma and urine samples from primary brain tumor patients using sequencing guided by mutations detected in multi-region tumor biopsies. When using this sensitive technique, we found no improvement in detection of tumor-derived DNA by analysing CSF samples. We also identified mutations in ctDNA obtained from plasma and urine during follow-up sampling post-surgery. We propose therefore that this method could also be used to track tumor recurrence.

## Results

In this study, we analysed a total of 73 samples including tumor subparts (3 to 6 per patient), CSF, plasma, buffy coat and urine, from 8 patients: 7 patients with primary glioblastomas and 1 patient with anaplastic oligodendroglioma (**Table S1** and **Table S2**). DNA from 34 tumor subparts from 8 biopsies were analysed by whole exome sequencing (WES), resulting in detection of 8838 single nucleotide variants (SNV) (**Fig. 1A**) at an average of 1105 SNV per patient (range 435-1725) (**Table S3** and **Fig. S1**). Mutations were called from the different tumor subparts individually, and also collectively by merging the sequencing reads from these subparts in order to increase the sensitivity for detecting low abundance stem-mutations of the common clones. Hybrid-capture sequencing panels were designed to target the SNVs detected by WES in the tumor (driver and passenger mutations, a total of 8838 loci) and this list was supplemented by comprehensive coverage of the 52 most frequently mutated genes in glioma (5). Sequencing data were analyzed using the INVAR (INtegration of VAriants Reads) pipeline, which combines locus-based noise filtering, strand selection, and enrichment of mutant fragments using ctDNA biological charcteristics (**Fig. S2, Fig. S3** and **Table S4**) (22). Fractions of tumor-derived DNA were estimated by integrated mutant allele fractions (IMAF) which indicate the fraction of reads covering target loci which carried the mutant alleles identified in the matched patient’s tumor specimens.

**Fig. 1:**
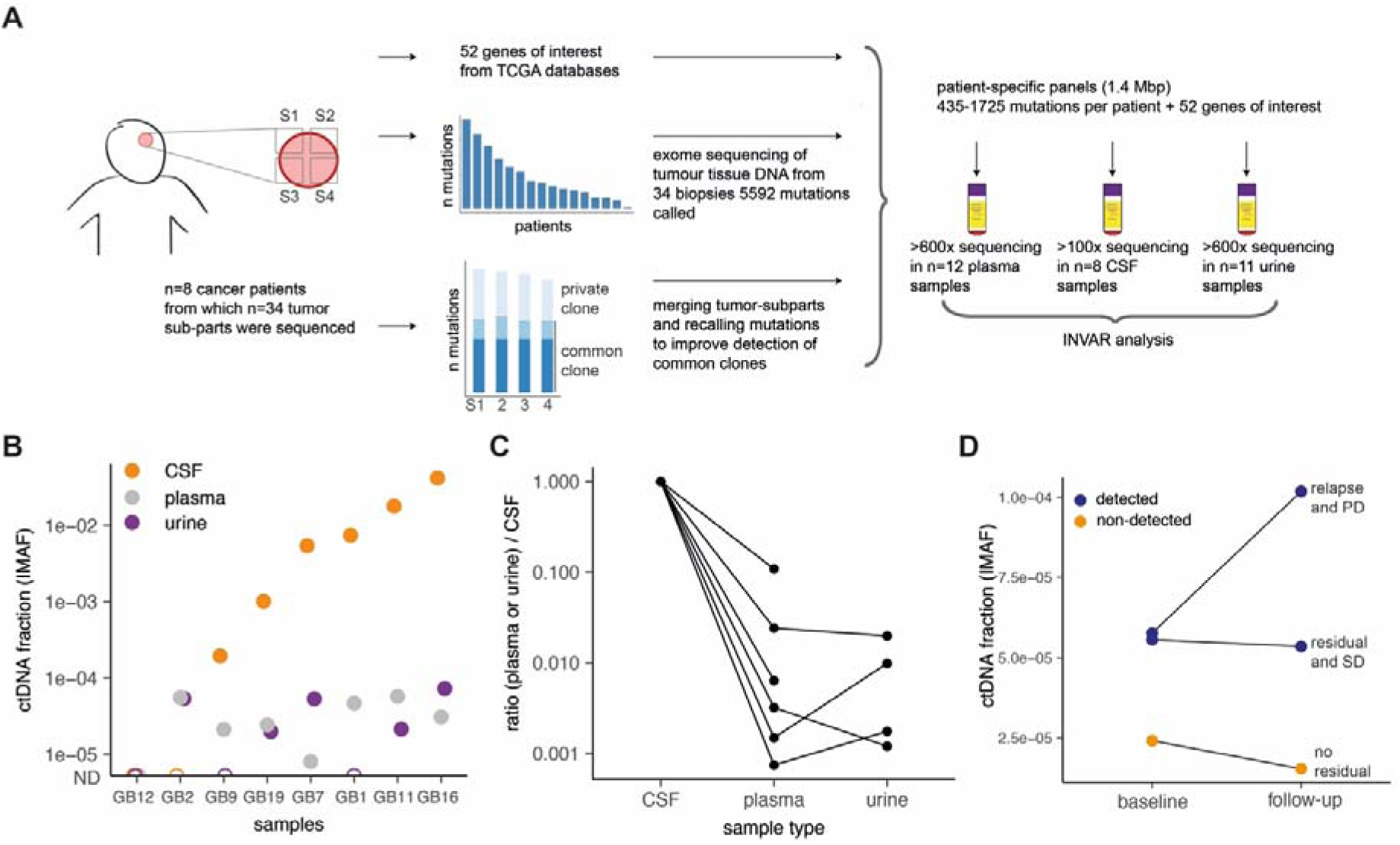
Integrating tumor clonality analysis improves the detection of ctDNA in CSF and plasma from gliomas patients. **A:** Schematic of ctDNA detection in matched CSF, plasma and urine samples from glioma patients using INVAR. Depth of sequencing indicated is the mean across the samples analyzed. **B:** Estimated ctDNA fractions for the matched plasma (7/8 detected), CSF (6/8 detected) and urine samples (5/8 detected) collected immediately pre-surgery from 7 patients with primary glioblastoma and one patient (GB12) with anaplastic oligodendroglioma. The ctDNA fraction is expressed as IMAF (Integrated Mutant Allele Fraction). Detected cases are indicated by full circle and non-detected cases as an open circle. ND: non-detected. **C:** Estimated relative ratio of tumor DNA fractions (IMAF) in plasma and urine compared to the matched CSF samples. One patient (GB12) had tumor DNA detected in plasma and urine but not in CSF and is not shown. **D:** Detection of residual and progressive disease in plasma samples taken within 6 months of treatment. Follow-up plasma samples were available for 3 patients (GB2, GB11 and GB12). Blue data points indicate the IMAF of samples with detected ctDNA GB11 with progressive disease, PD, and GB2 with stable disease, SD), whilst orange data points indicate the IMAF of samples (from GB12 with no residual disease observed) that were not confirmed as being detected (below the limit of calling). Data are annotated with clinical information (residual disease, relapse, PD and SD) determined by imaging.

ctDNA was detected in plasma from 7/8 patients prior to surgery, and in a total of 10/12 samples when including follow-up samples (**Fig. 1B**). ctDNA was detected in urine from 6/8 patients, in a total of 8/11 samples. Although more mutations were in general detected in CSF samples, when integrating data across all mutation loci there was no improvement in ctDNA detection rate by analysing CSF collected prior to surgery (6/8 patients detected). This was despite the mean ctDNA fraction being 243-fold lower in plasma (range 9-fold to 1343-fold) (**Fig. 1C**) and 389-fold lower in urine (range 50-fold to 834-fold) compared to CSF, with IMAF of 3.1×10^−5^ (plasma), 4.72 × 10^−5^ (urine), and 6.4×10^−3^ (CSF) (**Fig. 1B**).

For 3 patients, samples of both urine and plasma were obtained 6 months following surgery in addition to the baseline samples collected immediately prior to surgery. Mutated (tumor-derived) DNA was detected 6 months post-surgery in both plasma and urine samples for 2/3 patients (**Fig. 1D**). Contrast agent-enhanced T_1_-weighted MRI demonstrated that the two positive patients had residual or recurrent disease, whereas the patient with no detectable ctDNA had no evidence of recurrence. These preliminary data suggest that tumor-guided sequencing of urine or plasma samples may also be used to detect recurrent disease.

Of the 52 genes frequently mutated in glioma, mutations were detected at least once for the majority of genes across all tumor subparts (**Fig. 2A**). By limiting analysis in body fluids to these 52 most commonly mutated genes, which thus does not require pre-analysis of the tumor samples, we were able to detect mutations in plasma samples from 4/8 of cases, which was not improved by CSF analysis (also 4/8). Mutations in these genes were detected in the urine of 1/8 cases. Amongst the large number of tumor-specific mutations detected, several clinically actionable mutations were detected (**Fig. 2A** and **Fig. 2B**), Notably, *EGFR* and *MUC16* mutations were identified in plasma samples, both being potentially targetable with precision therapy (**Fig. 2A** and **Table S4**).

**Fig. 2:**
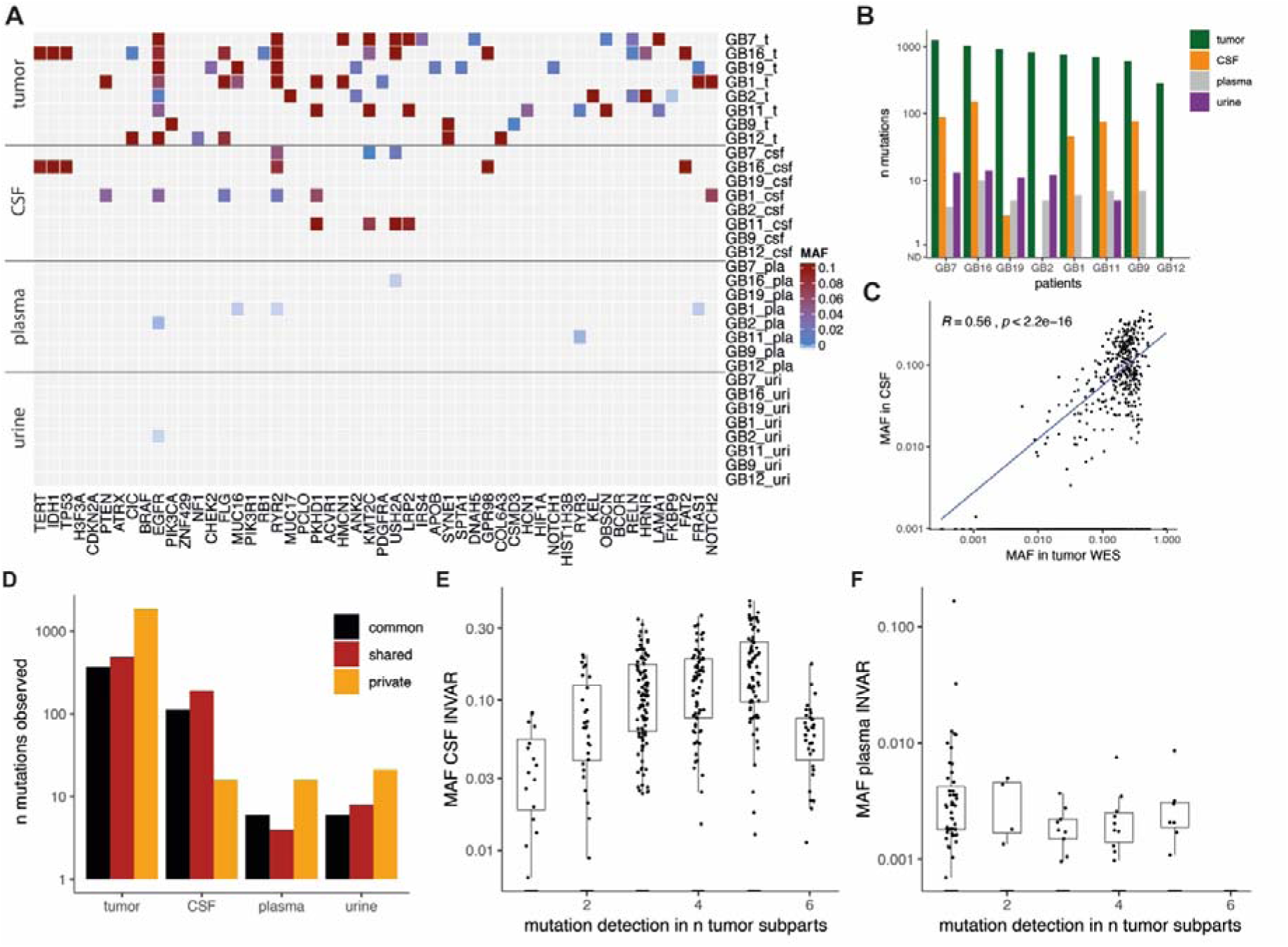
Detection of ctDNA in plasma, urine and CSF is affected by intra-tumor heterogeneity in gliomas. **A:** Mutant allele fraction (MAF) in tumor samples and body fluids for the detected mutations in the 52 most frequently mutated genes in gliomas based on the TCGA databases. **B:** Summary of the number of mutations detected in DNA from tumor tissue, CSF, plasma and urine samples in the baseline pre-surgery samples from the 8 patients. **C:** Comparison of the MAF of detected mutation in tumor tissue DNA by WES, with the MAF of the mutation detected in CSF by targeted sequencing. **D:** Mutations are classified depending on their respective clones (in black commonly detected in all tumor subparts, in red shared across some tumor sub-parts, and in orange private mutations). **E, F:** Mutations common and shared across multiples tumor tissue subparts are more frequently detected and with a higher MAF than private mutations in CSF samples (**E**), but not in plasma samples (**F**).

Detection of ctDNA in bio-fluids is thought to be affected by intra-tumoral genomic heterogeneity (23). The mutant allele fraction (MAF) in tumor DNA correlated with that of matched mutations detected in CSF (R=0.56, p<0.0001), but not in plasma (R=0.14, p>0.05) or urine (R=0.12, p>0.05) (**Fig. 2C** and **Fig. S4**). We subsequently compared the frequency of variant detection in plasma, urine and CSF samples with the presence of the mutation in one or more tumor sub-parts (**Fig. 2D**). Mutations common to all subparts, or that were shared across several sub-parts, were more frequently detected in CSF when compared with mutations that were private to a single region (**Fig. 2D**). This resulted in a trend for increased representation of clonal mutations in CSF samples (**Fig. 2E** and **Fig. S5**). However, in plasma and urine samples the opposite was observed where mutations tended to be representative of clones identified within a single tumor region (**Fig. 2F** and **Figure S6**).

## Discussion

We have shown that ctDNA is can be detected, with suitably sensitive methods, in the urine and plasma for the majority of patients with high grade glioma; and that analysis of CSF may not provide added benefit for detection of tumor-derived DNA when such methods are employed. ctDNA was detected in the urine (6/8) and plasma (7/8) of patients with glioma. By tracking a large number of mutations, we demonstrate that the sensitivity of detection can be improved. The median ctDNA fractions in plasma and urine were very low (IMAF of 3.1×10^−5^ and 4.72×10^−5^ respectively).

Whilst methylation-based detection, cell-free DNA genome-wide fragmentation, tumor-derived mitochondrial DNA (mtDNA), exosomes, vesicles and tumor-educated platelets have all been proposed as alternate methods for plasma-based detection of glioma-derived mutant DNA, these alternate strategies provide limited information about the tumor genome (7, 24–28). Through tumor-guided sequencing we have identified mutations in the plasma and urine of GB patients, which has the potential to track tumor-specific actionable mutations that may be important for targeting therapies and for monitoring tumor recurrence.

In other cancers, ctDNA has been shown to be representative of treatment-driven clonal tumor evolution, with real time measurements of clonal dynamics reflecting clonal tumor hierarchy (23, 29). The use of multi-region tumor sampling allowed us to identify clonal, shared and private mutations within the primary cancer (30), and on the matched CSF, plasma and urine samples. We show that the representation of these mutations varies between the plasma and urine samples, which mostly represented private clones, and the CSF which mostly representated clonal or shared mutations.

Overall, we demonstrated that ctDNA can be detected in the plasma and urine of patients with GB, a cancer that has previously been beyond the reach of non-invasive liquid biopsy.

## Materials and Methods

### Study design

Patients were recruited at Addenbrooke’s Hospital, Cambridge, UK as part of the BLiNG (Biopsy of Liquids in New Gliomas) study (REC reference number: 15/EE/0094) (Supplementary Table 1). Matched tumor tissue DNA, CSF, plasma, urine and buffy coat samples were collected for each patient. Written informed consent was obtained from the patients; the studies were conducted in accordance with the Declaration of Helsinki and were approved by an Institutional Review Board. Lumbar puncture was performed immediately prior to craniotomy for tumor debulking. After sterile field preparation, the thecal sac was cannulated between the L3 and L5 intervertebral spaces using a 0.61 mm gauge lumbar puncture needle, and 10 ml of CSF was removed. After collection, CSF and whole blood samples were immediately placed on ice and then rapidly transferred to a pre-chilled centrifuge for processing. Samples were centrifuged at 1500 g at 4C for 10 minutes. Supernatant was removed and further centrifuged at 20,000 g for 10 minutes, and aliquoted into 2 mL microtubes for storage at −80 °C (Sarstedt, Germany).

### Sample preparation

Tumor tissue DNA were extracted and isolated as described previously (15). Fluids were extracted using the QIAsymphony platform (Qiagen, Germany). Up to 10 mL of plasma, 10 mL of urine and 8 mL of CSF was used per sample. DNA from plasma, urine and CSF samples was eluted in 90 µL, and further concentrated down to 30 µL using a Speed-Vac concentrator (Eppendorf, Germany).

### Sequencing library preparation and WES for tissue DNA

In order to identify patient specific somatic mutations, we first performed whole exome sequencing (WES) of all tumor tissue and germline buffy coat DNA samples. Fifty nanograms of DNA were fragmented to ∼120bp by acoustic shearing (Covaris) according to the manufacturer’s instructions. Libraries were prepared using the Thruplex DNA-Seq protocol (Rubicon Genomics) with 5x cycles of PCR. Libraries were quantified using quantitative PCR (KAPA library quantification, KAPA biosystems) and pooled for exome capture (TruSeq Exome Enrichment Kit, Illumina). Exome capture was performed with the addition of i5 and i7 specific blockers (IDT) during the hybridization steps to prevent adaptor ‘daisy chaining’. Pools were concentrated using a SpeedVac vacuum concentrator (Eppendorf, Germany). After capture, 8x cycles of PCR were performed. Enriched libraries were quantified using quantitative PCR (KAPA library quantification, KAPA Biosystems), DNA fragment sizes were assessed by Bioanalyzer (2100 Bioanalyzer, Agilent Genomics) and captures were pooled in equimolar ratio for paired-end next generation sequencing on a HiSeq4000 (Illumina).

Sequencing reads were de-multiplexed, allowing zero mismatches in barcodes. The reference genome was the GRCh37/b37/hg19 human reference genome - 1000 Genomes GRCh37-derived reference genome, which includes chromosomal plus unlocalized and unplaced contigs, the rCRS mitochondrial sequence (AC:NC_012920), Human herpesvirus 4 type 1 (AC:NC_007605) and decoy sequence derived from HuRef, Human Bac and Fosmid clones and NA12878. The sequence data of the patient samples were aligned to the reference genome using BWA-MEM v0.7.15. Duplicate reads were marked using Picard v1.122 (http://broadinstitute.github.io/picard). Somatic SNV and indel mutations were called using GATK Mutect2 (Genome Analysis Toolkit), (https://www.broadinstitute.org/gatk) in tumor-normal pair mode using buffy coat as the normal.

MAFs for each single-base locus were calculated with MuTect2 for all bases with PHRED quality ≥30. After MuTect2, we applied filtering parameters so that a mutation was called if no mutant reads for an allele were observed in germline DNA at a locus that was covered at least 10x, and if at least 4 reads supporting the mutant were found in the plasma data with at least 1 read on each strand (forward and reverse). Variants were annotated using Ensembl Variant Effect Predictor with details about consequence on protein coding, accession numbers for known variants and associated allele frequencies from the 1000 Genomes project.

### Tumor-guided capture sequencing

Hybrid-based capture for the different body fluids (CSF, plasma, urine) analysis was designed to cover the variants identified above for each patient using the SureDesign software (Agilent). In addition, 52 genes of interest for glioma were included in the tumor-guided sequencing panel based on the TCGA databases. Patients were separated into 2 panels covering all the mutations included for these patients (4 patients per panel). Patients GB1, GB2, GB9, GB16 were grouped in panel 1, and patients GB7, GB11, GB12, GB19 were grouped in panel 2. Panel 1 covers in total 526 kbp (5841 regions) and panel 2 covers 526 kbp as well (5701 regions). Custom panels ranged in size between 1.404-1.430 Mb with 120 bp RNA baits. Baits were designed with 5x tiling density, moderately stringent masking and balanced boosting. 99.7% of the targets had baits designed successfully.

Indexed sequencing libraries were prepared using the Thruplex tag-seq kits (Takara). Libraries were captured either in 1-plex for plasma and urine samples or 3-plex for CSF samples (to a total of 1000 ng capture input) using the Agilent SureSelectXTHS protocol, with the addition of i5 and i7 blocking oligos (IDT), as recommended by the manufacturer for compatibility with ThruPLEX libraries. Custom Agilent SureSelectXTHS baits were used. 13 cycles were used for amplification of the captured libraries. Post-capture libraries were purified with AMPure XT beads, then quantified using quantitative PCR (KAPA library quantification, KAPA Biosystems), and DNA fragment sizes controlled by Bioanalyzer (2100 Bioanalyzer, Agilent Genomics). Capture libraries were then pooled in equimolar ratios for paired-end next generation sequencing on a HiSeq4000 (Illumina).

### Capture sequencing analysis and INVAR

Sequencing reads were de-multiplexed, allowing zero mismatches in barcodes. Cutadapt v1.9.1 was used to remove known 5’ and 3’ adaptor sequences specified in a separate FASTA of adaptor sequences. Trimmed FASTQ files were aligned to the UCSC hg19 genome using BWA-mem v0.7.13 with a seed length of 19. Error-suppression was carried out on ThruPLEX Tag-seq library BAM files using CONNOR. The consensus frequency threshold -f was set as 0.9 (90%), and the minimum family size threshold -s was varied between 2 and 5 for characterization of error rates. For capture data, a minimum family size of 2 was used.

Patient-specific sequencing data consists of informative reads at multiple known patient-specific loci that were identified from tumor sequencing (see above). Because each panel comprised mutations from multiple patients, we can compare mutant allele fractions across loci as a means of error-suppression. Patients could control for another patient’s panel as long as the tumor sequencing did not identify any overlapping mutations. The distribution of signal across loci potentially allows for the identification of noisy loci not consistent with the overall signal distribution. Loci that carried signal in more than 10% of control samples or a mean allele fraction >1% were blacklisted as noisy and removed from the analysis. Each locus was also annotated with trinucleotide error rate, the corresponding tumor allele fraction, fragment size and whether that locus passes an additional outlier suppression filter as identified by INVAR (INtegration of VAriant Reads), (version 0.7).

For each sample, the IMAF (Integrated Mutant Allelic Fraction) was determined across all loci passing pre-INVAR data processing filters with mutant allele fraction at that locus of <0.25; loci with signal >0.25 mutant allele fraction were not included in the calculation because (i) loci would not be expected to have such high mutant allele fractions in body fluids of glioma patients (unless they are mis-genotyped SNPs), and (ii) if the true IMAF of a sample is >0.25, when a large number of loci are tested, they will show a distribution of allele fractions such that detection is still supported by having many low allele fraction loci with signal. Based on the ctDNA level of the sample, the binomial probability of observing each individual locus given the IMAF of that sample was calculated. Loci with a Bonferroni corrected P-value <0.05 (corrected for the number of loci interrogated) were excluded in that sample, thereby suppressing outliers.

## Supplementary Materials

Fig. S1. Quality control assessment of the WES data from the tumor tissue DNA.

Fig. S2. Quality control and background sequencing error rates of the two capture sequencing panels.

Fig. S3. Mutant and non-mutant DNA fragment size distribution determined from the capture sequencing data.

Fig. S4. The number and MAF of detected mutations increase with the number of tumor sub-parts in CSF.

Fig. S5. Matched mutations detected and intra-tumor heterogeneity for each patient depending on the type of samples.

Fig. S6. MAF of the detected mutations depending on the tumor subparts and bio-fluids.

Table S1. Characteristics of the patients included in the study.

Table S2. Characteristics of the samples included in the study.

Table S3. List of SNVs called by Mutect2 after WES of the tumor tissue DNA individual sub-parts.

Table S4. Patient-specific variants passing filters after tumor-guided sequencing of the CSF, plasma and urine samples.

## Acknowledgments

We wish to thank for their help and support the Cancer Research UK Cambridge Institute core facilities, in particular bio-repository, bioinformatics and genomics. We wish also to thank the Cambridge Molecular Diagnostic Laboratory.

## Author contributions

Concept and design of the study: F.M., N.R. and R.M.; Methodology: F.M., K.H., J.C.M.W, C.G.S. and R.M.; Investigation: F.M., K.H., M.T., C.G.S., D.G., I.H., W.C., M.G., J.C.M.W., I.H., W.C., D.G. and R.M.; Sample Collection: R.M. and C.W.; Data Analysis: F.M., K.H., N.R. and R.M.; Computational analysis: F.M., J.S., C.G.S., J.M., J.C.M.W., M.E., D.C. and K.H.; Writing – Original Draft: F.M. and R.M.; Writing – Review & Editing: F.M., R.M., K.H., I.H., C.G.S, C.W., K.B., and N.R.; Funding Acquisition: C.W., K.B., N.R., R.M.; Supervision: F.M., K.B., N.R. and R.M.

## Data and materials availability

Liquid biopsy sequencing data will be made available at the European Genome-phenome archive. The INVAR pipeline will be made publicly accessible at http://www.bitbucket.org/nrlab/invar.

