## Supplementary_material for "Integrated clonal analysis reveals circulating tumor DNA in urine and plasma of glioma patients"

^§^ Current affiliation: Precision Medicine, R&D Oncology Unit, AstraZeneca, CB4 0WG, Cambridge, UK.

^†^ Current affiliation: Institute of Cancer Genomics Science, University of Birmingham, B15 2TT, Birmingham, UK.

**Fig. S1:**

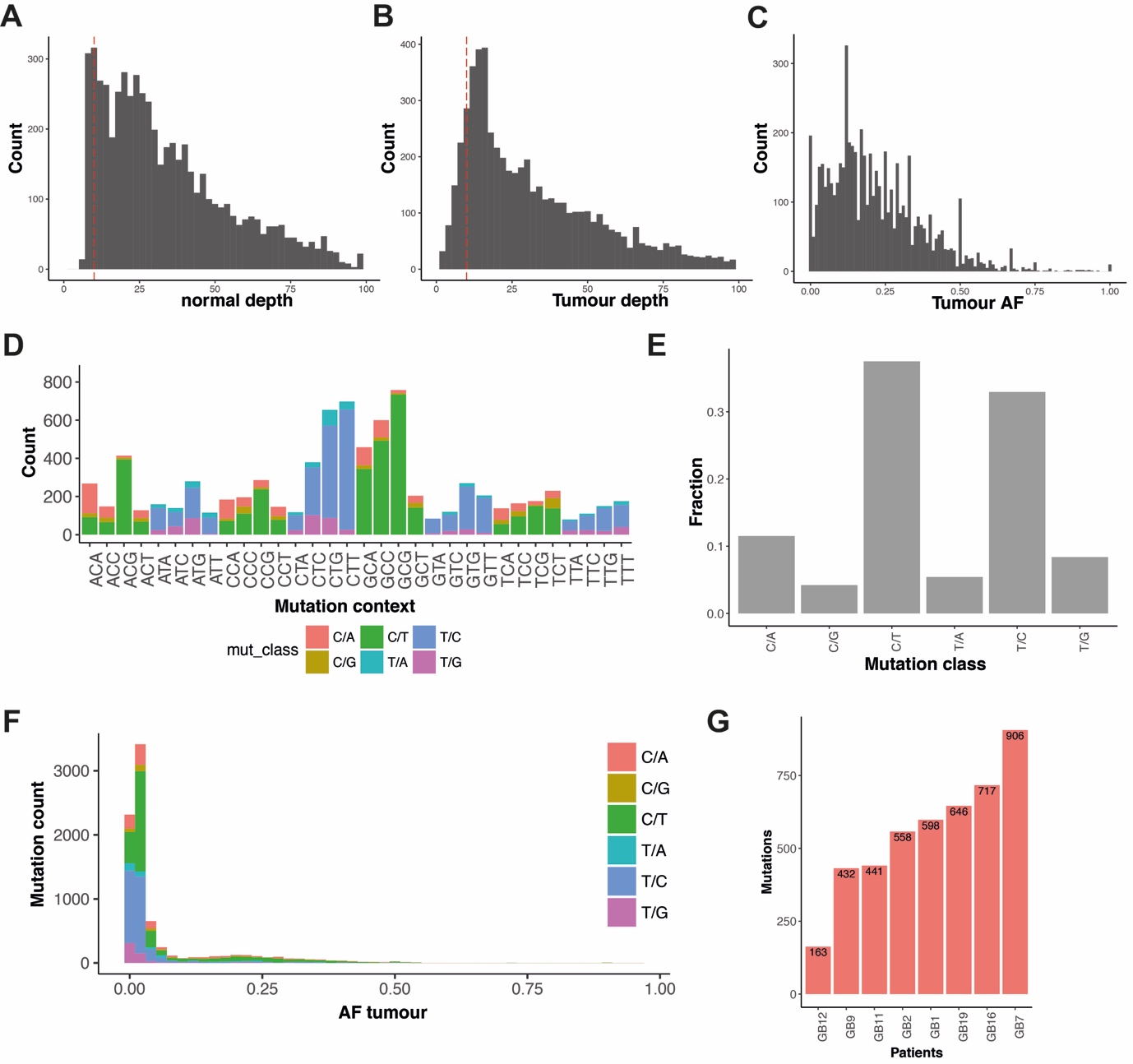

**Fig. S1: quality control assessment of the WES data from the tumor tissue DNA.** **A:** Sequencing depth from white blood cell DNA extracted from the buffy coat layer. **B:** Sequencing depth from the tumor tissue DNA. **C:** Mutant allele fraction (MAF) for the variants called from WES of the tumor tissue. **D:** Mutation counts by trinucleotide context, colored by mutation class. **E:** Frequency of each mutation class included in the capture panels. **F:** Distribution of tumor mutation allele fractions in tumor samples colored by mutation class after individual and merged variant calling. **G:** Number of mutations per patient retained after filtering.

**Fig. S2:**

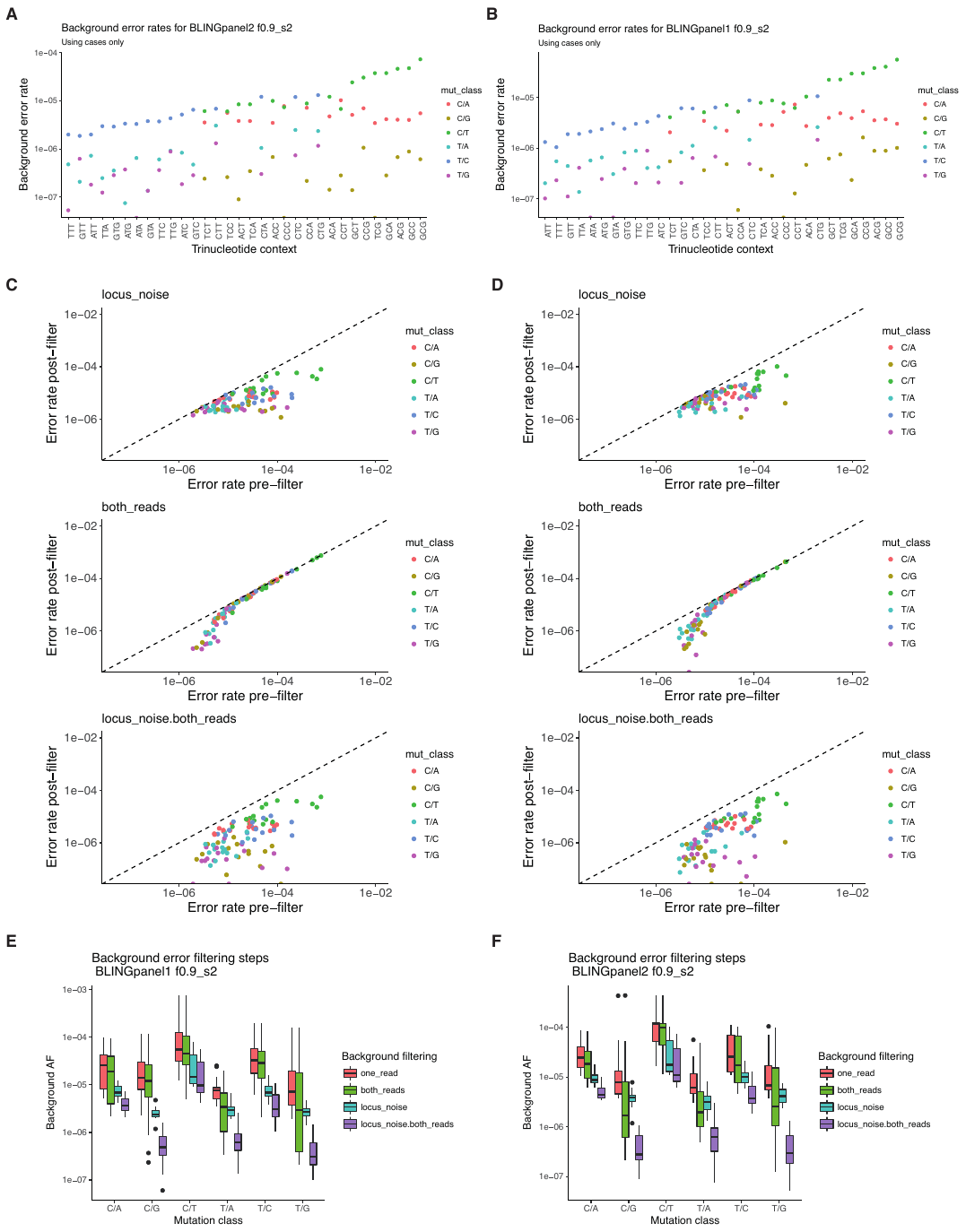

**Fig. S2: quality control and background sequencing error rates of the two capture sequencing panels.** Sequencing error rates were calculated from plasma samples. **A:** Background error rates per mutation class for the first capture sequencing panel. **B:** Background error rates per mutation class for the second capture sequencing panel. **C:** Background error rates per trinucleotide for the first capture sequencing panel are plotted before and after each background error filtering steps for each trinucleotide context. **D:** Background error rates per trinucleotide for the second capture sequencing panel are plotted before and after each background error filtering steps for each trinucleotide context. **E:** Summary of the error rate per class depending on the filtering steps applied for the first capture sequencing panel. **F:** Summary of the error rate per class depending on the filtering steps applied for the second capture sequencing panel.

**Fig. S3:**

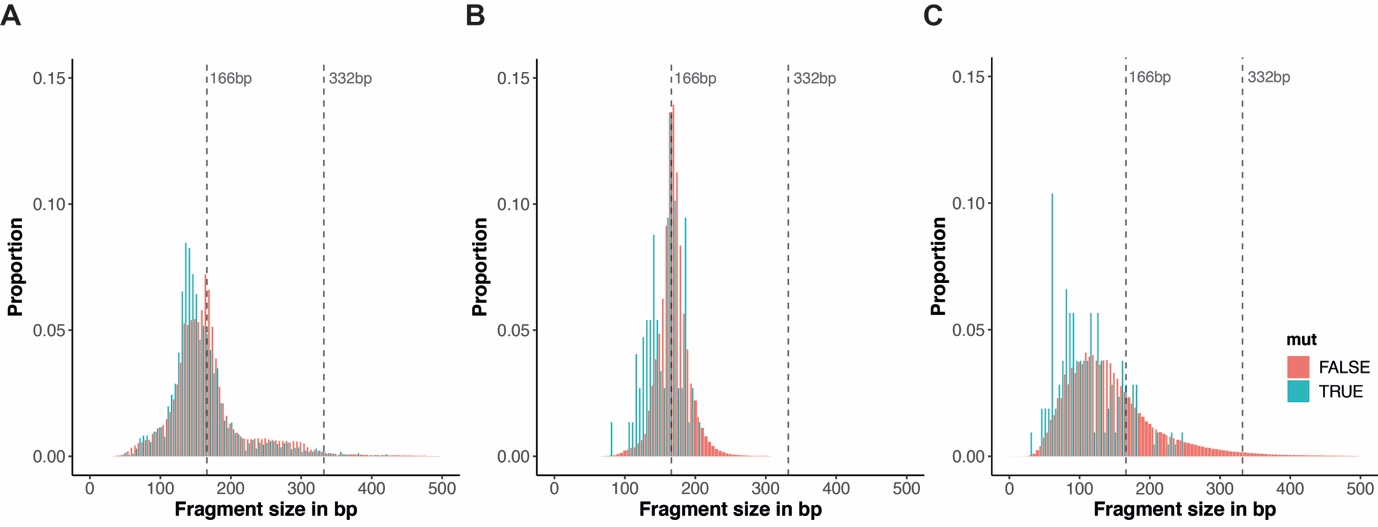

**Fig. S3: mutant and non-mutant DNA fragment size distribution determined from the capture sequencing data.** Shown in red are the size distribution of non-mutant reads and in blue the size distribution of mutant reads. **A:** Size distribution from the merged reads of the CSF samples. **B:** Size distribution from the merged reads from the plasma samples. **C:** Size distribution from the merged reads from the urine samples.

**Fig. S4:**

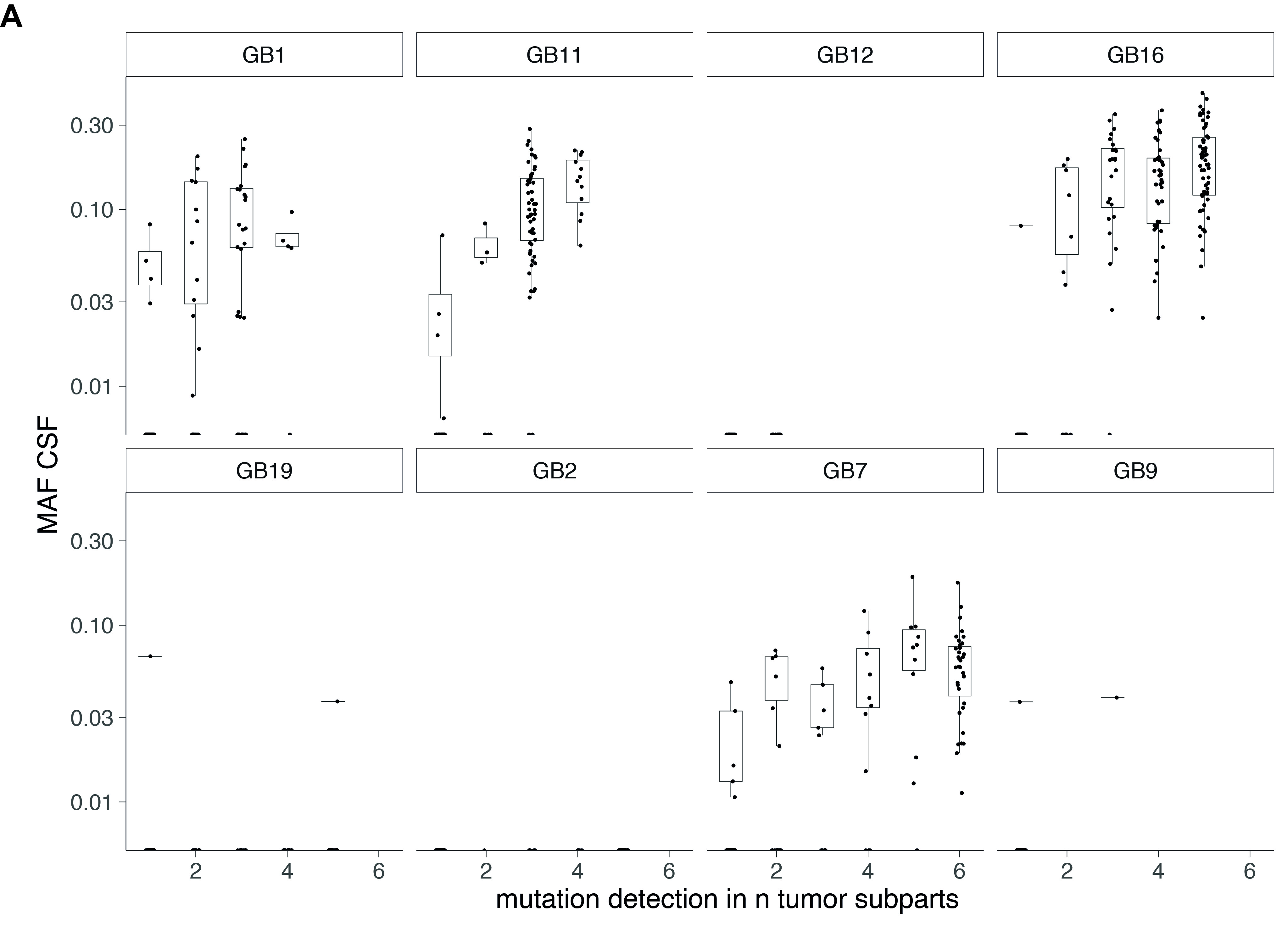

**Fig. S4: the number and MAF of detected mutations increase with the number of tumor sub-parts in CSF.** MAF were determined for each mutation detected in the CSF samples. The mutations are distributed depending on their frequency of detection per tumor-subparts.

**Fig. S5:**

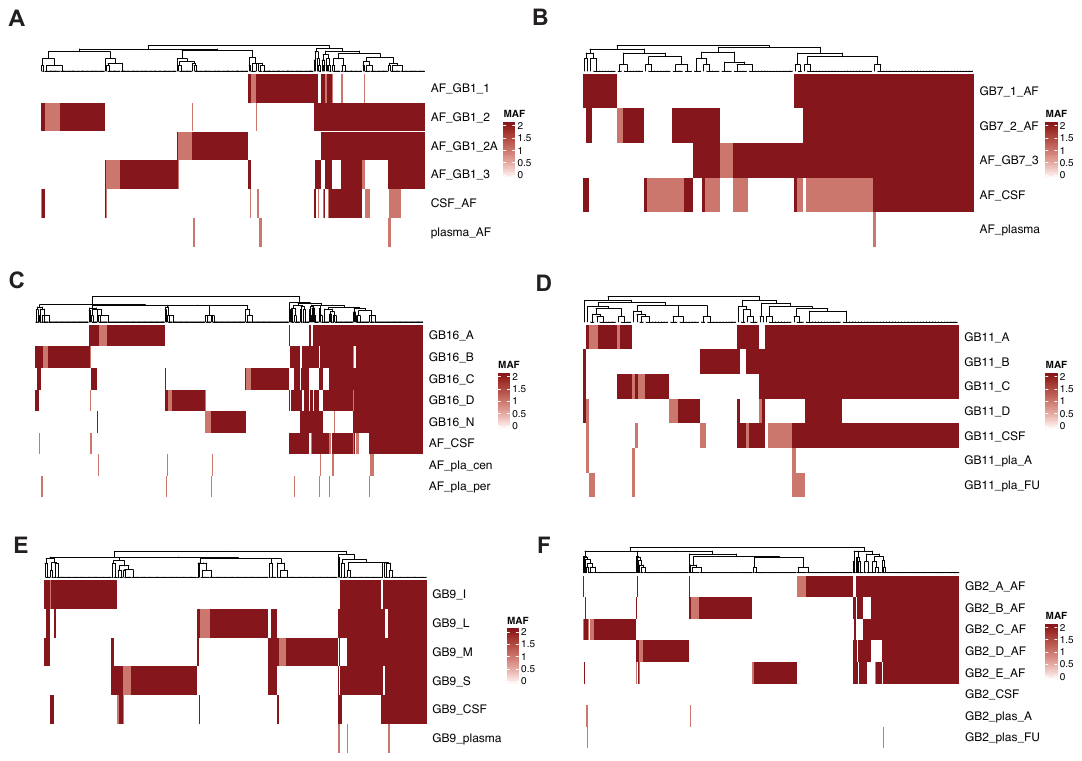

**Fig. S5: matched mutations detected and intra-tumor heterogeneity for each patient depending on the type of samples.** Shown in red are the MAF for the mutations detected in the tumor tissue DNA, CSF, plasma and urine for each patient. When available, follow-up plasma samples were included. For the tumor tissue DNA, details of the different tumor sub-parts are provided. **A:** Patient GB1; **B:** Patient GB7; **C:** Patient GB16; **D:** Patient GB11; **E:** Patient GB9; **F:** Patient GB2.

**Fig. S6:**

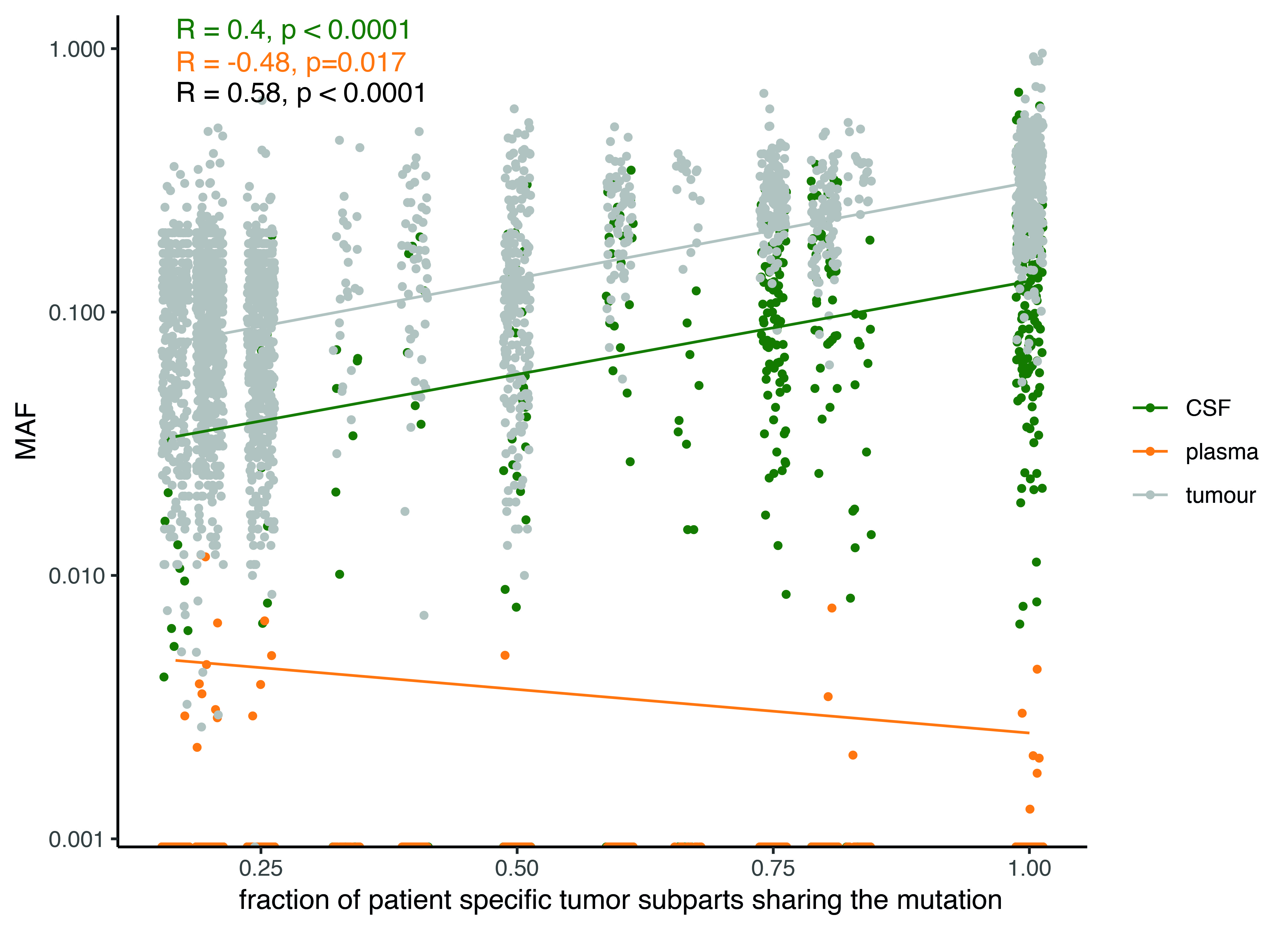

**Fig. S6: MAF of the detected mutations depending on the tumor subparts and bio-fluids.** MAF were determined for each mutation detected in the tumor tissue DNA, CSF and plasma samples for all patients using INVAR. MAF is plotted against the fraction of patient specific tumor subparts sharing that mutation (the fraction of subparts is a discrete variable with values which are fractions with integer values ranging from 1 to 6 for both nominator and denominator; jitter is added to improve data visualization).

**Table S1:** Characteristics of the patients included in the study.

**Table S2:** Characteristics of the samples included in the study.

**Table S3:** List of SNVs called by Mutect2 after WES of the tumor tissue DNA individual sub-parts.

**Table S4:** Patient-specific variants passing filters after tumor-guided sequencing of the CSF, plasma and urine samples.
